# Psychophysics with R: The R Package MixedPsy

**DOI:** 10.1101/2022.06.20.496855

**Authors:** Priscilla Balestrucci, Marc O. Ernst, Alessandro Moscatelli

## Abstract

Psychophysical methods are widely used in neuroscience to investigate the quantitative relation between a physical property of the world and its perceptual representation provided by the senses. Recent studies introduced the Generalized Linear Mixed Model (GLMM) to fit the responses of multiple participants in psychophysical experiments. Another approach (two-level approach) requires fitting psychometric functions to each individual participant data using a Generalized Linear Model (GLM), and then testing the hypotheses on the multiple participants by means of a second level analysis. For either options, the implementation of the statistical analysis in R is possible and beneficial. Here, we introduce the package **MixedPsy** to model and fit psychometric data in R, either with two-level and GLMM approaches. The package, freely available in the CRAN repository, uses different methods for the estimation of Point of Subjective Equivalence (PSE) and Just Noticeable Difference (JND), and provides utilities for immediate visualization and plotting of the fitted results. This manuscript aims to provide researchers with a practical tutorial for implementing a complete analysis pipeline for psychophysical data using **MixedPsy** and other packages and basic functionalities of the R programming environment.

## Introduction

Psychophysics investigates the quantitative relation between a physical property of the environment and its perceptual representation provided by the senses. For example, researchers may be interested in investigating the perceived brightness of a visual stimulus, the speed of an object sliding across the skin, or the perceived pitch of a sound. The relation between stimulus intensity and perception can provide evidence about fundamental sensory processes, assess the performance characteristics of an observer, and provide indications for the necessary specifics of a technological device (Pelli and Bart 1991). Psychophysics has a long history of methodological research, where different experimental protocols and statistical models have been proposed. A popular and powerful experimental method consists in forced-choice tasks, where participants are asked to judge a given feature of the stimulus by selecting one among a discrete set of possible responses. The relationship between the intensity of the physical stimulus and the probability of the response in this type of task is modeled at the participant level by the psychometric function estimated with Generalized Linear Models (GLM) (Agresti 2002; Knoblauch and Maloney 2012). In our previous study, we showed that it is also possible to use Generalized Linear *Mixed* Models (GLMM) to estimate the responses of multiple participants at the population level (Moscatelli, Mezzetti, and Lacquaniti 2012).

Several publications over the last decade have illustrated the advantages of using R for fitting GLMs and GLMMs to psychophysical data (Yssaad-Fesselier and Knoblauch 2006; Knoblauch and Maloney 2012; Moscatelli, Mezzetti, and Lacquaniti 2012; Linares, López-Moliner, et al. 2016). R is a modular, open source software environment and programming language for statistical computing and graphics, that compiles and runs on Linux, (Mac) OS X, and Windows platforms (R Core Team 2020). It is also one of the most common tools for statistics and data science, and among the 20 most popular programming languages in 2022, according to the [TIOBE index](https://www.tiobe.com/tiobe-index/).

To facilitate modeling and visualization of psychophysical data in R, we have recently developed the open source package **[MixedPsy](https://cran.r-project.org/package=MixedPsy)** (Moscatelli and Balestrucci 2021). In this tutorial, we will show how to use our package together with other R resources to process, analyze, and communicate results of psychophysics experiments. While a sound grasp of the relevant model frameworks is a prerequisite for a full understanding of the proposed R syntax, as well as for correctly interpreting the analysis outcome, this tutorial is meant for researchers irrespective of their programming background or experience with R. Please note that this is not meant to be a tutorial on mixed models: For such a tutorial, refer to (Agresti 2002; Moscatelli, Mezzetti, and Lacquaniti 2012; Knoblauch and Maloney 2012). For a comprehensive and detailed overview on linear mixed modeling and GLMM fitting, refer to (Bates et al. 2015) by authors and maintainers of package **lme4**.

This tutorial is organized as follows. First, we will shortly describe an experimental procedure typically used to collect data in psychophysics, and the models most commonly applied to this type of data: the psychometric function and the Generalized Linear Mixed Model (GLMM). Next, we will provide R code examples of the syntax needed for fitting psychometric functions and GLMMs, using **stats** and **lme4** packages, respectively. Throughout the tutorial, we will provide examples of how to manipulate and process data using packages from the popular **tidyverse** suite (Wickham and Grolemund 2016). While such processing can be done in many different ways, including basic R scripting, we chose to show this approach as we feel that it provides a compact solution, easy to read and especially beneficial for those with a limited programming background.

### Experimental Methods in Psychophysics

The forced-choice task is a common experimental procedure in psychophysics (Rohde, Dam, and Ernst 2016). This procedure allows a bias-free measurement of discrimination performance if it is not possible to systematically identify one stimulus from the other. With this method, participants are presented with two stimuli that differ in a continuous property (e.g., their size, timing, location, etc.) and have to perform a discrimination judgment (e.g. which of the two stimuli they perceive as larger/shorter/closer than the other). In forced-choice tasks, one of the two stimuli is a reference and the other is a test stimulus, whose intensity varies across different trials of the experiment (Kingdom and Prins 2016). Each intensity level of the test stimulus is repeated several times in randomized order throughout the experiment, so that it is possible to evaluate the performance on the psychophysical task, given by the proportion of correct responses for each intensity level. Throughout this tutorial, we will consider two different dataset resulting from forced-choice tasks, both of which are included in the **MixedPsy** package and described in detail in subsequent sections.

The relation between intensity of a physical stimulus and behavioral performance is modeled by fitting a psychometric function. The psychometric function allows to determine quantitative parameters that summarize the performance of an observer, and evaluate the underlying sensory mechanisms involved. Two key parameters are the Point of Subjective Equality (PSE) and the Just Noticeable Difference (JND). The PSE indicates the intensity level in which a stimulus is perceptually equivalent to another of different magnitude. JND gives the minimum difference in intensity required discriminate between two stimuli for a specified percentage of trials. Typically and unless otherwise specified, such percentage is fixed at 50%. In other words, PSE and JND provide a measure for the accuracy and precision of the response, respectively (Kingdom and Prins 2016).

### Analysis of the binary response of individual participants: The GLM

The psychometric function is a special application of the Generalized Linear Model (GLM) (Agresti 2002; Moscatelli, Mezzetti, and Lacquaniti 2012; Knoblauch and Maloney 2012) and relates the intensity of the presented stimulus to the probability of the binary response (e.g., “comparison longer” or “comparison shorter” in a forced-choice task judging the length of a physical stimulus, etc.). The model equation is the following:

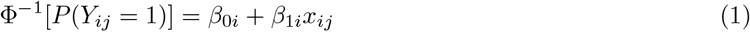

The response variable *P*(*Y_ij_* = 1) is the probability, for participant *i* and observation *j*, of responding 1 or 0 (mapped respectively as *longer* and *shorter, yes* and *no*, etc.) given the stimulus level *x_ij_* of the predictor variable *x*. The probit link function Φ^−1^(·) allows to establish, between the response probability and the continuous predictor, a linear relationship, which is fully described by two parameters: intercept *β*_0_ and slope *β*_1_. Other link functions like Logit and Weibull are also often used in psychophysics (Agresti 2002; Klein 2001). More complex models have been proposed to take into account lapses and guesses by the participant (Wichmann and Hill 2001). Similarly, in detection tasks (“yes” or “no” responses) the guessing parameter sets the lower asymptote of the curve at 0.5, with the the midpoint of the curve at 0.75 being the detection threshold. These models will not be covered in this tutorial.

Parameters relative to observer *i*, PSE_i_ and JND_i_, are computed from intercept and slope in Eq.1 as follows:

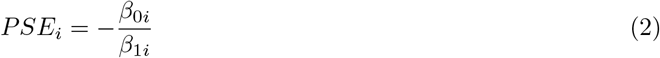

and

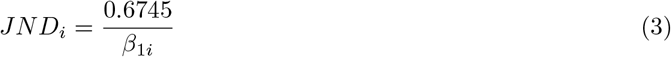

where 0.6745 is the value of the 75th percentile of the normal distribution. Often, when using a probit function, the JND is calculated at the 84th percentile (i.e. one standard deviation of the normal distribution), which corresponds to 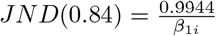.

In a typical psychophysics experiment, not only do we want to estimate PSE and JND associated with manipulation of a continuous stimulus (such as time, velocity, weight, etc.), but we also want to assess how these parameters are modulated by some factorial or categorical predictor (e.g. gender, handedness, presence or absence of a confounding stimulus, etc.). To do this, the model in Eq. 1 must be applied to each observer and for each experimental condition, and the parameters of the resulting psychometric functions (Eq. 2–3) are used as inputs for the second-level analysis (Moscatelli, Mezzetti, and Lacquaniti 2012), for example by means of t-tests or analysis of variance (ANOVA).

### The analysis of the response at the population level: The GLMM

By definition, a GLMM is an extension to clustered data of a GLM. In a psychophysics experiment, the data are “naturally” divided in clusters, since multiple responses are collected from each observer participating to the experiment. Intuitively, the responses from a single participant will tend to be more correlated with each other than with the responses from other participants. GLMMs include this information when evaluating the relation between the probability of the response and the predictors, by simultaneously fitting both the effects associated with the experimental variables and the differences necessarily occurring between participants (Agresti 2002). The model equation is formalized as follows:

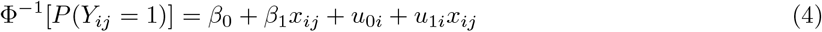

Parameters *β*_0_ and *β*_1_, akin to the parameters of the psychometric function in Eq. 1, estimate the effects of the experimental variables on the probability of the response, and are called *fixed effects*. Parameters *u*_0*i*_ and *u*_1*i*_ estimate the heterogeneity between individual participants in the population sample (i.e. how intercept and slope change for each participant *i*), and are called *random effects*. For GLMMs, we assume that the random-effect parameters are normally distributed random variables.

Advantages of the GLMM with respect to the two-level analysis are the clear distinction between the between- and the within-participants variability and the higher statistical power (Moscatelli, Mezzetti, and Lacquaniti 2012).

Given the model in Eq. 4, we can obtain PSE and JND using the same relationship described in Eq. 2 and 3. In this case, however, the estimates refer to the fixed-effects at the sample level, and not to the parameters associated to the performance of a single participant.

The GLMM in Eq. 4 is a univariable model, because it evaluates the relationship between the probability of the response and one continuous predictor. As such, it is the extension at the population level of the psychometric function in Eq. 1. As already noted, in psychophysics we typically not only want to measure the response to a certain stimulus, but also evaluate whether and how such response changes under different conditions. While this further comparison requires a second-level analysis when fitting GLMs to individual participants’ responses, it can be evaluated within the same model with multivariable GLMMs (Moscatelli, Mezzetti, and Lacquaniti 2012; Hidalgo and Goodman 2013). In fact, the GLMM equation can then be further extended to account for effects associated with multiple predictors:

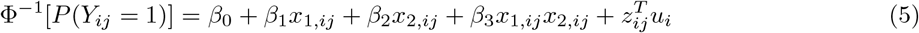

In Eq. 5, the model includes parameters for two predictors, *x*_1_ and *x*_2_, as well as for their interaction; the term 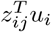 includes the associated random effects. Such a model can very easily represent a common situation in psychophysics, where we vary the intensity of a physical stimulus, represented by a continuous variable *x*_1_, under two (or more) experimental conditions, which correspond to the levels of a categorical variable *x*_2_.

In principle, a model can include many coefficients associated with different fixed and random effects. As a general rule, it is recommended to avoid fitting data with an overly complicated model, for several reasons: First, a model with less coefficients is easier to interpret. Second, adding free parameters may lead to data overfitting. This is a phenomenon that occurs when the model follows not only the signal, but also the noise associated with the data, too closely. Overfitting should be avoided, because it may decrease the ability of the model to generalize or predict the response from a different sample of the same population the dataset comes from. In this sense, we can think of the dataset as training data for our model, regardless of whether or not we have a “test” dataset available (James et al. 2013).

Deciding which are the important predictors to include in the model is a fundamental step in the analysis pipeline with a GLMM. Such selection typically requires fitting models including different subsets of predictors, inspect the model residuals, and compare them based on some statistics such as the Akaike Information Criterion (AIC) or the Bayesian Information Criterion (BIC). Note that, as the number of possible predictors increases, comparing all possible models becomes quickly unfeasible, and it is necessary to restrict the search for the best fitting model to a subset of possible models, e.g. subset selection, forward or backward stepwise selection, or dimension reduction (James et al. 2013). Discussing these different approaches for model selection is out of the scope of this tutorial, and interested readers can refer to (James et al. 2013).

### Variance estimation of the PSE, JND

When fitting a GLM or GLMM with the related function in R,`stats::glm()` and `lme4:glmer()` respectively, we automatically obtain the estimated parameters and their standard error, i.e. the uncertainty associated with the estimate. PSE and JND are calculated as non-linear combinations of the parameters of the GLM/GLMM (i.e., the intercept and the slope in a univariable model). Therefore, the variance of the PSE and JND cannot be estimated directly from the standard error of the model parameters. In our previous article (Moscatelli, Mezzetti, and Lacquaniti 2012) we showed how to easily estimate PSE and JND and their associated variance using either the delta method or the bootstrap method, and these routines are included in **MixedPsy**. The delta method is based on the assumption that the estimator is asymptotically approximated to a normal distribution, and calculates mean and variance of a function of random variables by using its truncated Taylor series (Casella and Berger 2002). While this method is relatively straightforward and computationally inexpensive, the assumption of asymptotic normal distribution is not necessarily correct for GLMMs. With the non-parametric bootstrap method, the variance of the estimator is obtained by re-sampling with replacement from the original dataset. The bootstrap samples are then used to compute the estimates of the parameters of interest and their variance (James et al. 2013). The bootstrap approach does not make any assumption regarding the sample distribution, therefore it is suitable for evaluating estimates from GLMM parameters. The accuracy of the estimation relies on the high number of bootstrap samples. As a consequence, this procedure can be computationally costly.

## Methods

### The R environment

This tutorial has the scope to provide the necessary information to get started using relevant R packages, such as **stats**, **lme4**, and **MixedPsy**, for the analysis of psychophysical data. We encourage the reader to have an R environment running and the necessary packages installed and loaded, in order to replicate the examples proposed in the next sections. For more information and resources on the R software and the packages used in this section, refer to the Appendix.

### Setup

The R code described in this tutorial was tested on an Apple MacBook Pro computer running under macOS 10.16 and on a Dell Latitude computer running Ubuntu Linux 20.04. The R version was 4.0.2 (released 2020-06-22), with loaded packages **MixedPsy** (1.1.0), **lme4** (1.1-27.1), **tidyverse** (1.3.1). For a detailed report of the environment used, refer to the Appendix.

### Dataset

**MixedPsy** includes two different psychophysical datasets that will be used as benchmarks throughout the tutorial.

`simul_data` is a simulated dataset created using the **MixedPsy** function `PsySimulate()`. It simulates the results that would be obtained from an experiment where observers are presented with two stimuli of different intensity (e.g. duration), and have to judge which of the two is greater than the other. The independent variable `X` is the intensity of the stimulus and has nine levels, equally spaced and ranging between 40 and 120 (arbitrary units). The binomial response is summarized in the `Longer` and `Total` columns, representing how many times the simulated subjects judged the test stimulus as more intense (i.e. longer) than the reference stimulus, and the number of repeated measures for each intensity level, respectively. Invoking the **MixedPsy** function `PsySimulate()` without changing the default settings generates a dataset analogous to simul_data. Since `PsySimulate()` is based on random numbers generation, the values of associated with each stimulus level will not be exactly the same. If for reproducibility issues, one wants to obtain the same random numbers as those in the package dataset, define `set.seed(123)`. For details on how to simulate custom data, refer to the **MixedPsy** package documentation.

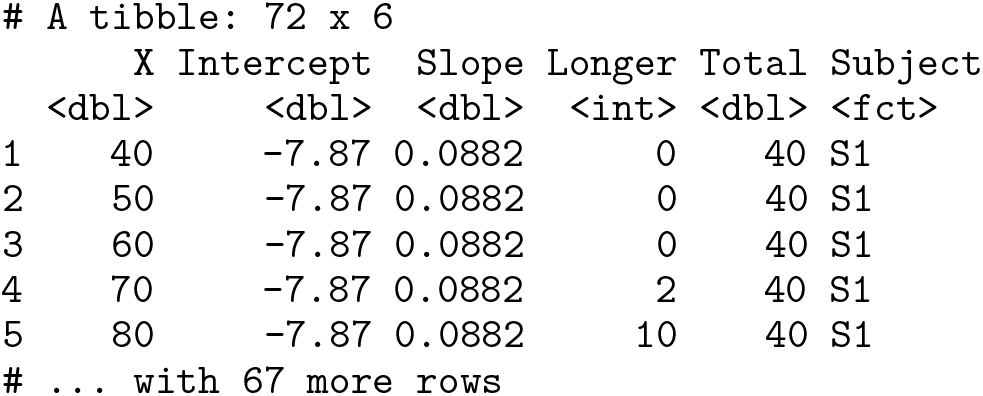

The second dataset, `vibro_exp3`, was collected from real observers in a psychophysics study published in (Dallmann, Ernst, and Moscatelli 2015). In the experiment, nine participants were required to judge the motion speed of a moving surface by touching it, and decide whether it was faster or slower than a reference stimulus, while exposed to different vibratory noise conditions. The speed of the test stimulus was chosen among seven speeds ranging between 1.0 and 16.0 cm/s, and the masking vibrations had a frequency of either 0 or 32 Hz (i.e., control and test condition, respectively). Each combination speed/vibration was repeated 40 times in randomized order, resulting in a total of 560 trials. The dataset is organized in 5 columns, each representing one of the variables of interest, namely: speed of the test stimulus (`speed`), frequency of the masking vibration (`vibration`), repetitions in which the subject perceived the stimulus as faster than the reference, repetitions judged as slower (`faster` and `slower`, respectively), observer ID (`subject`).

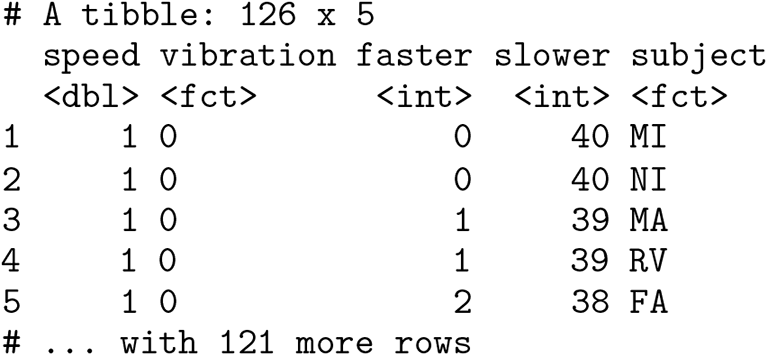

Typically, the output file of a psychophysics experiment is a data frame where every row correspond to a single observation, but note that the functions in **MixedPsy** cannot process input data in this form. Instead, the data frame must have a summarized structure, where each row in the table contains all the information related to one level of the independent variables under consideration (i.e. observer ID, stimulus intensity, experimental conditions, etc.). While the data provided in **MixedPsy** already have this structure, a preliminary processing step must be carried out when importing data from a performed experiment. This can be done using functions of the **tidyverse** suite. A representative raw file can be downloaded from the materials included in the tutorial, and imported in the R environment with `read_csv()`:

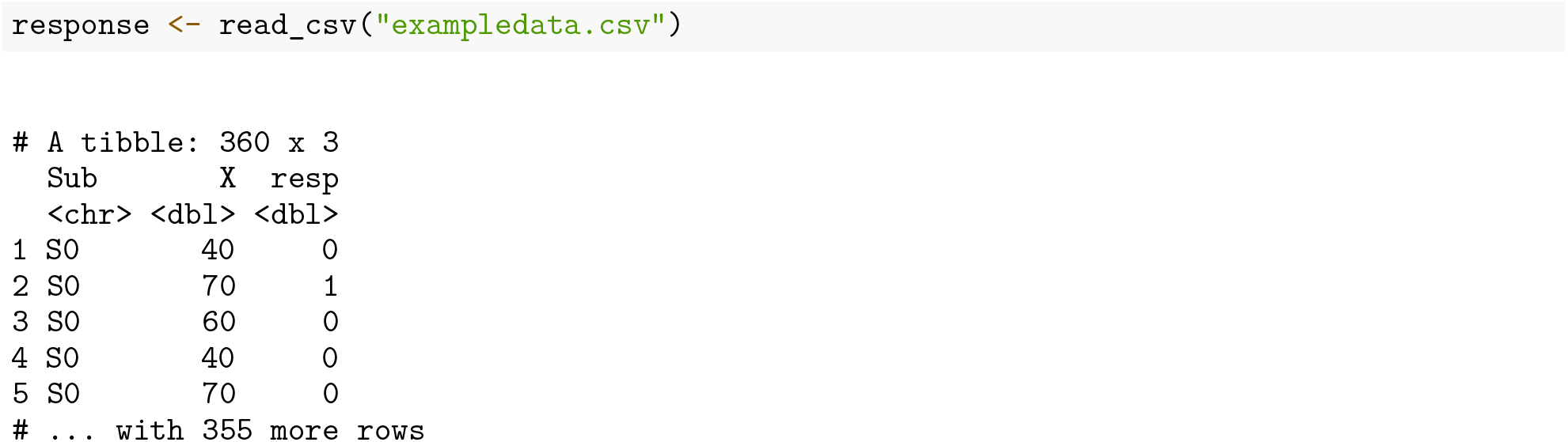

To obtain a data frame that can be used in further analysis, we need to collapse all the observations in relation to a given intensity level into a single row, and compute for each intensity the sum of responses (counts) that were mapped with level 1 and those mapped to 0. With functions in the tidyverse, the code is the following:

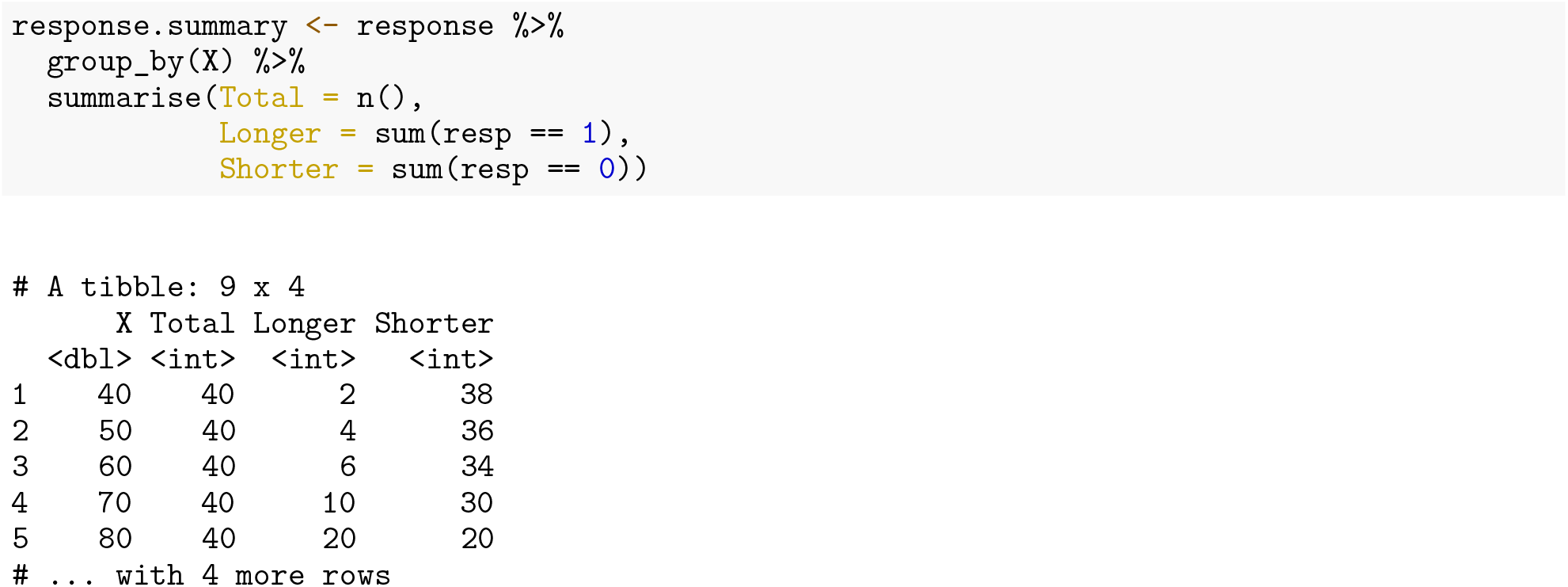

What the code does is straightforward: a new object is defined by taking the imported dataset, grouping it by levels of the `X` variable, then summarizing for each level the total number of responses, as well as which were mapped as 0 and 1. The pipe operator `%>%` allows to pass intermediate results from one function to the next without having to explicitly define and store them, and makes the code concise and easy to read. An intuitive way to understand its function is to pronounce it as “then” when reading the code (Wickham and Grolemund 2016).

## Results

### Fitting GLMs with R: The Psychometric Function

In this section, we describe how to fit a psychometric function to a set of example data. To fit a GLM to the response of one of the simulated observers in `simul_data`, we need to extract all the rows for Subject `S1` from the data frame:

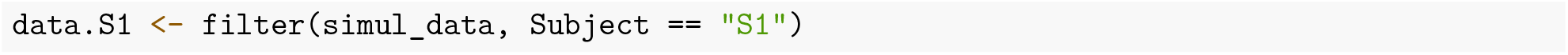

Next, we can fit a GLM to the response of `S1` using `glm()`, which is part of the `stats` package and therefore loaded by default in any R environment, as follows:

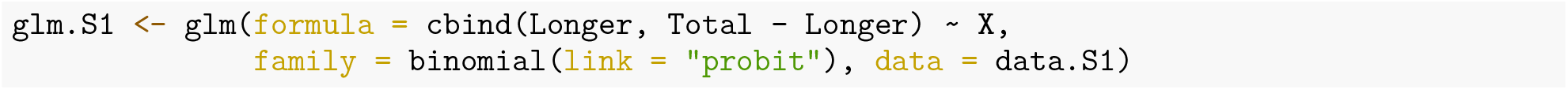

In accordance with the definition in Eq. 1, the function takes the relation between response and predictor variable as input (the term before and after the tilde ~ symbol, respectively) in the `formula` argument. The binomial response variable is usually specified as a two-column matrix, with each column providing the number of successes and failures, respectively. The two columns are combined together with function `cbind()`. Unless otherwise specified, the intercept of the model is included by default and not explicitly defined in the formula. The `family` argument contains information about the class of the error distribution and about the `link` function (`binomial` and `probit`, respectively). Other possible link functions that can be specified in `glm()` for binomial data are `logit`, `cauchit`, `log`, and `cloglog`.

A summary of the model is available with command `summary(glm.S1)`:

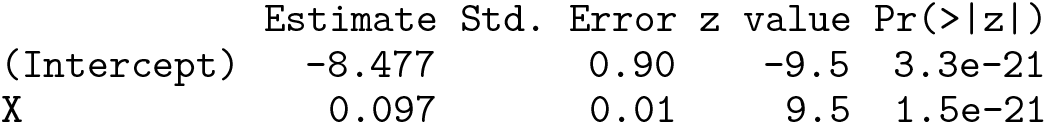

The first and second column show estimate and standard error of intercept and slope of the linear predictor function, which correspond to coefficients *β*_0*i*_ and *β*_1*i*_ of Equation 1, respectively. The last two columns report z-score and significance level of the test for the null hypothesis **H_0_:** *β* = 0 (Agresti 2002). In the example, both intercept and slope are significantly different from 0, and the null hypothesis can be rejected.

To estimate PSE and JND of the psychometric function for observer `S1` (Eq. 2 and 3), we use function `PsychDelta()` from **MixedPsy**. This calculates the parameters from a fitted GLM with a probit link function, and estimates their variance using the Delta method (Casella and Berger 2002):

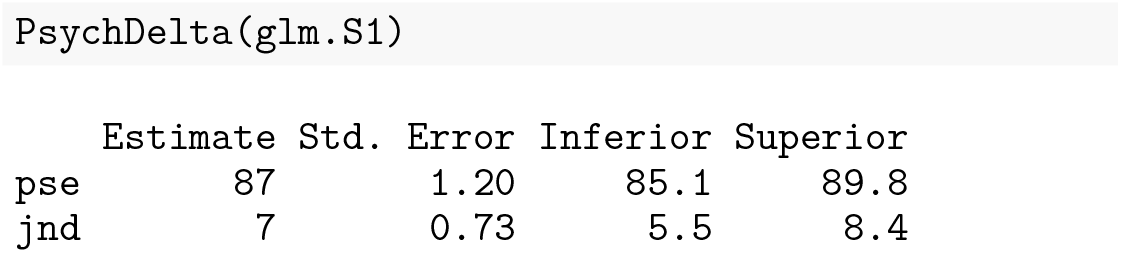

Finally, we can plot the psychometric function from the fitted GLM using `PsychPlot()`.

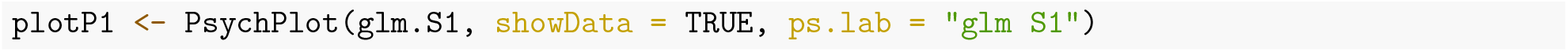

The output of the code above is shown in Fig. 1. By default, the function creates a new `ggplot` object. Alternatively, the the psychometric function can be added on an existing plot specified in argument `addTo`.

**Figure 1:**
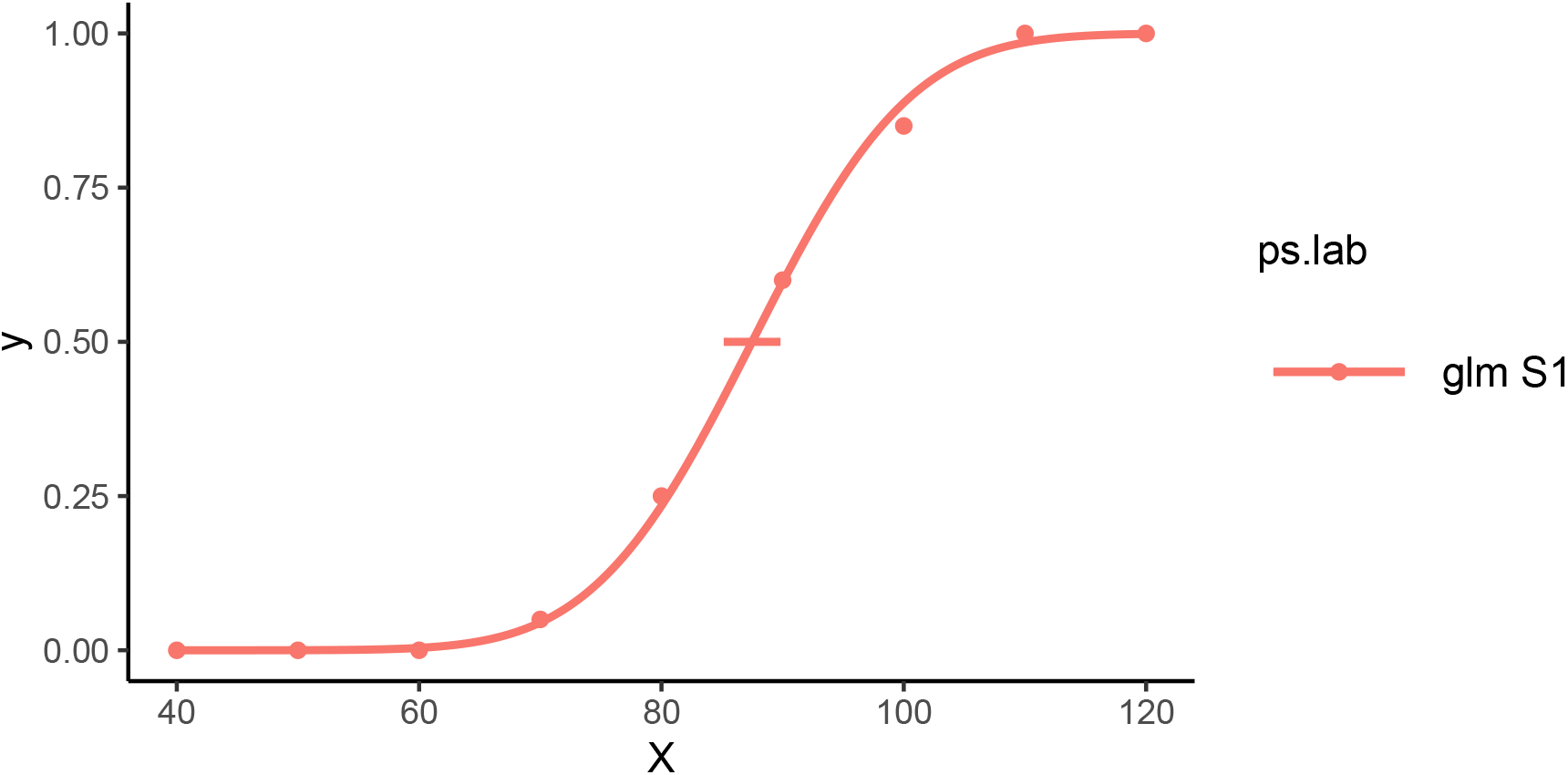
Visualization of the participant response and fitted psychometric function with `PsychPlot()`.

### Second-level analysis

In a typical experiment, the processing pipeline described above must be repeated for all observers and –if applicable– for all experimental conditions. PSE and JND for all observers must then be arranged in a data frame object for subsequent analysis (exploratory data analysis, inferential statistics, etc.). Here, we show a way to implement the routine described for a single participant (i.e. fitting the model with `glm()`, extracting PSE and JND with `PsychDelta()`) for all participants in our simulated dataset with an iterative process.

First, we define a function that contains the processing routine:

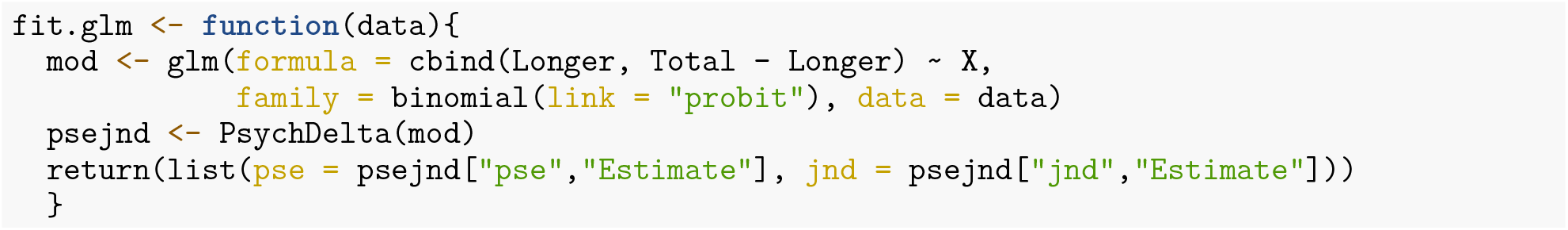

This function takes a data frame containing our variables of interest as input, fits a GLM to the data, and returns a `list` object containing the estimates of PSE and JND. We want to call this function for each subset of `simul_data` for a single participant, and the functions in the tidyverse suite provide a compact way of doing so:

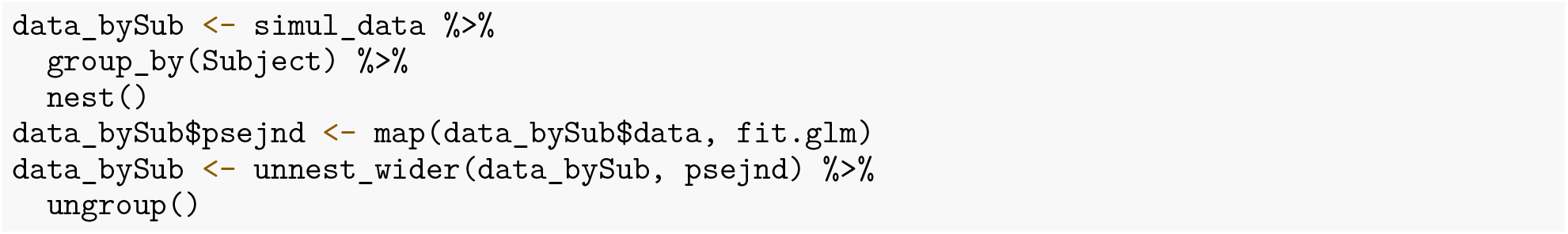

The `nest()` function rearranged the original data frame so that the `data` column contains a list of data frames grouped by `Subject`. Once we created the nested dataset, we created a new column (`psejnd`) by applying the previously defined function `fit.glm()` to every row of the `data` column, using the `map()` function. Finally, we assigned a separate column to each element of the lists in `psejnd` with function `unnest_wider()`, and removed the initial grouping operation with `ungroup()`.

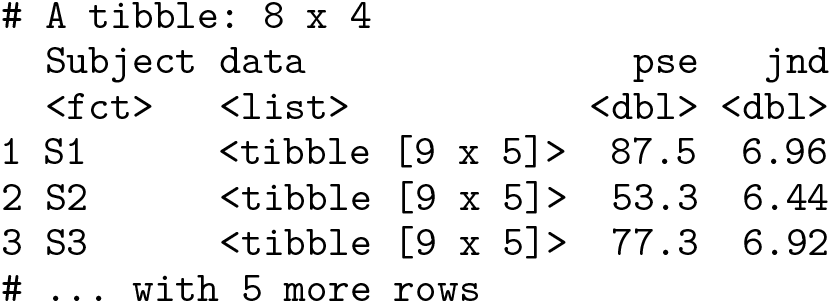

Given the list of PSE and JND for all participants in our dataset, we can get descriptive information of our sample by calculating measures of location and spread, as well as test hypotheses on the population with inferential statistics. For example, we can test whether the PSE is different from the mean of the presented stimulus intensities (in this case `X = 80`) by means of a one-sample t-test:

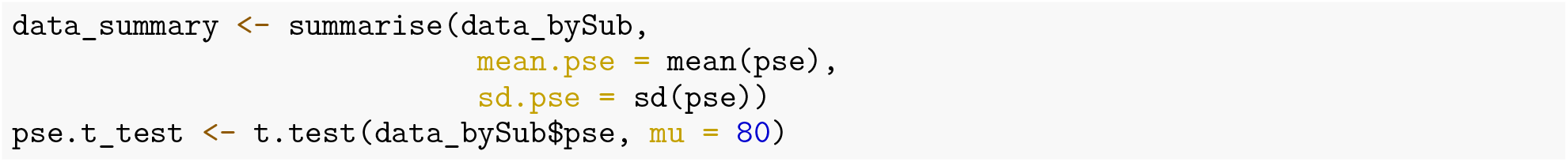

From this test, it emerges that the PSE is close to the mean value of the range of tested stimulus intensities (M = 75.4, SD = 14.32), and the null hypothesis cannot be rejected (t(7) = −0.91, p = 0.39).

In the next sections, we will show how to use packages **lme4** and **MixedPsy** to fit a GLMM and estimate PSE, JND, and their respective confidence intervals, in the case of a model with only a continuous predictor (univariable model), and with a continuous and a factorial predictor (multivariable model).

### Fitting GLMMs with R: Univariable case

For the univariable example, we will fit a GLMM as the one in Eq. 4 to the same simulated dataset used above with function `glmer()` from package **lme4**.

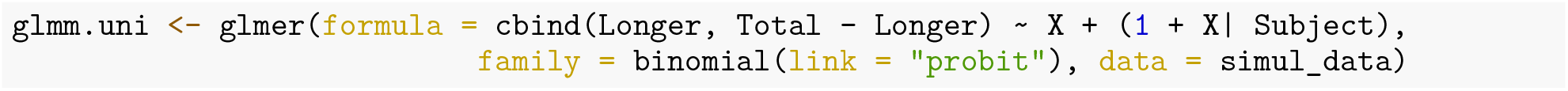

The syntax for `glmer()` is similar to that for `glm()`: we specify the relation between variable and linear combination of the predictors in the `formula` argument, and define the family of the error distribution and the link function in the `family` argument. The difference between `glm()` and `glmer()` lies in the way predictors are defined in the formula: in `glmer()`, the parameters outside parentheses account for the fixed effects (*β_i_* in Eq. 4), those in parentheses represent the random effects (*u_ij_* in Eq. 4). The clustering factor, in this case `Subject`, is declared in parenthesis, after the | symbol.

In the example above, we included intercepts and slopes as random effects in the model, meaning that both parameters change randomly across participants. However, we can evaluate whether the response can be described with a simpler model where the response of different participants changed only for the intercept value but not for the slope—that is, the psychometric functions of different participants are shifted one to the other, but they have all the same slope. In other words, they have different PSE but same JND. The model with only random intercept and no random slope is the following:

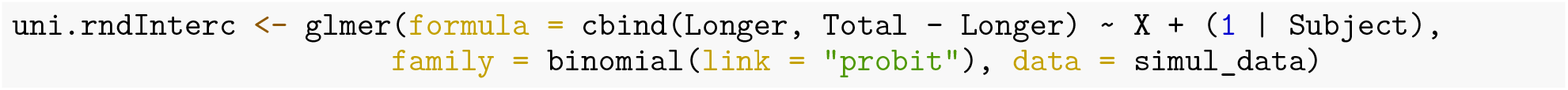

In order to decide which of the two models introduced above should be chosen for fitting the data, we can compare their AIC scores with the R function `anova()`:

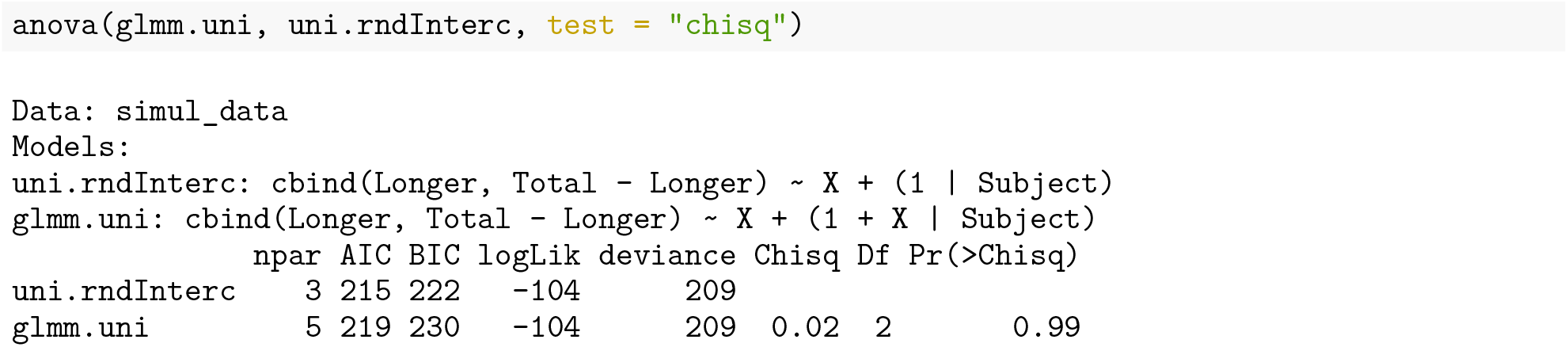

The AIC is based on a trade-off between goodness of fit and model complexity, so that the preferred model is the one with the lowest score (Akaike 1973). In the example, the AIC score favors the model with only random intercept (AIC = 215) over the one with both random effects (AIC = 219), and the likelihood ratio test does not provide evidence for the more complex model (p = 0.99). In such cases, the preference is always given to the model with less free parameters (i.e. smaller `npar`, model `uni.rndInterc` in our example.)

Once we have chosen the model, we use **MixedPsy** functions to plot the response, and to estimate PSE and JND of the population sample. For an easier maintenance of our package, most **MixedPsy** functions providing utilities that depend on package **lme4** do not take the model objects of class `glmerMod` as input. Instead, an object of class `xplode` must be created in an intermediate step:

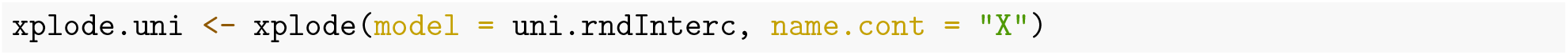

In order to define the object correctly, `explode()` takes the model object as well as the name of the continuous predictor as input, and –if present– the name of the factorial predictor.

Given the `xplode` object, we can visualize the response and relative model fit with `MixPlot()` (Fig. 2):

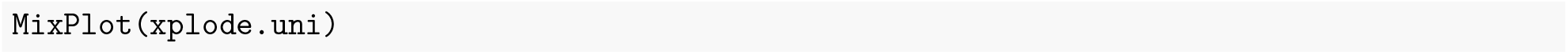

**Figure 2:**
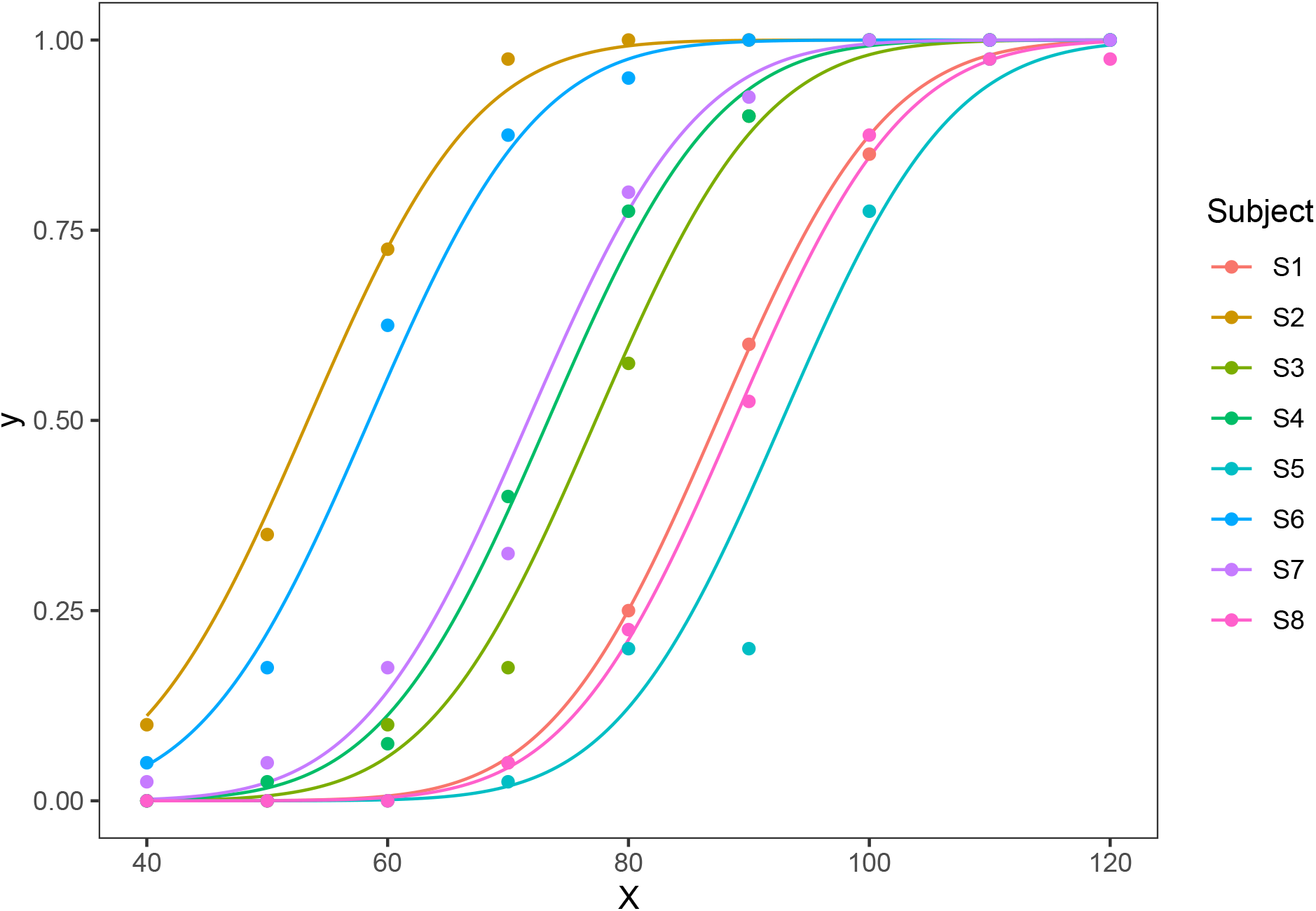
Plot wit MixPlot. GLMM predictions and point data.

`MixDelta()` provides estimate and 95% confidence intervals of PSE and JND evaluated with the delta method:

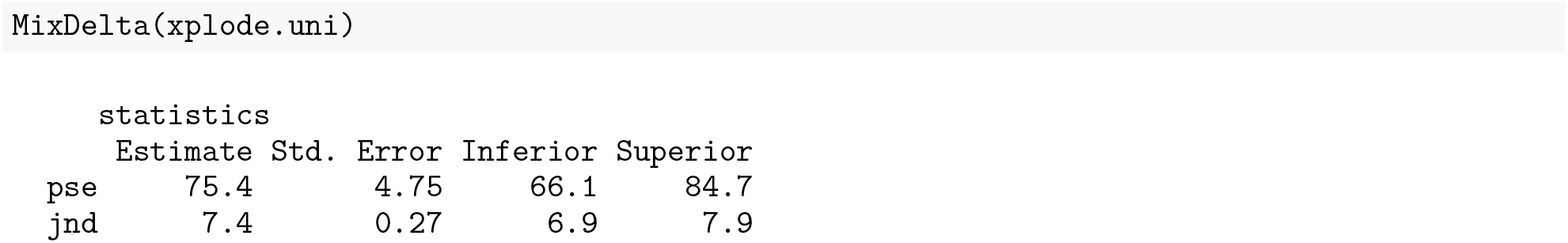

As mentioned in previous paragraphs, the delta method is based on asymptotic assumptions of a normal distribution of the parameters, which are not necessarily met in GLMMs. For a more correct estimate, `pseMer()` applies a bootstrap procedure on the object of class `glmerMod` and with a number of bootstrap samples specified in the `B` argument. For a univariable GLMM, `pseMer` provides estimates of the sample’s PSE and JND by default. Note that increasing the number of iterations will make the estimate more reliable, while also increasing the time needed for computation. With our setup, the estimation reported in this example with 500 iterations took 36.14 seconds to be completed.

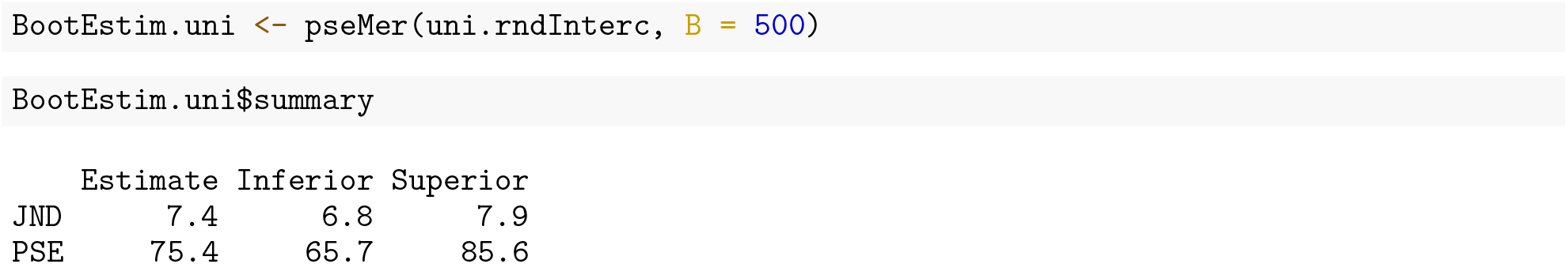

When comparing parameter estimations with the two different methods, we notice a slight difference in the confidence intervals, but not in the mean estimation. However, in both cases the confidence interval for the PSE includes the mean of the tested stimulus interval `X = 80`. This result is in accordance with that of the t-test introduced above for PSE values calculated with GLMs.

### Fitting GLMMs with R: Multivariable case

Multivariable GLMMs can be extremely useful to fit psychophysical data and test our experimental hypotheses. In this paragraph, we introduce the R syntax for fitting a GLMM with one continuous and one categorical predictor, and estimate PSE and JND associated with each level of the categorical predictor. To carry out the example, we use to the `vibro_exp3` dataset included in **MixedPsy**. For brevity, we do not discuss issues related to goodness of fit here. Note, however, that the analysis for model selection shown for predictors of the univariable GLMM applies also for multivariable GLMMs.

The `glmer()` syntax corresponding to the formula in Eq. 5 is the following:

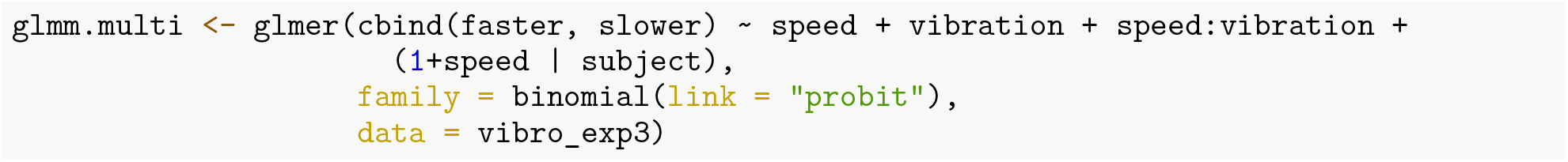

The fixed intercept corresponding to coefficient *β*_0_ is implicit. The same fixed effect parameters used in the model can be defined in an equivalent but more concise notation by including two independent predictors divided by the * symbol.

Let us look at the model’s coefficients:

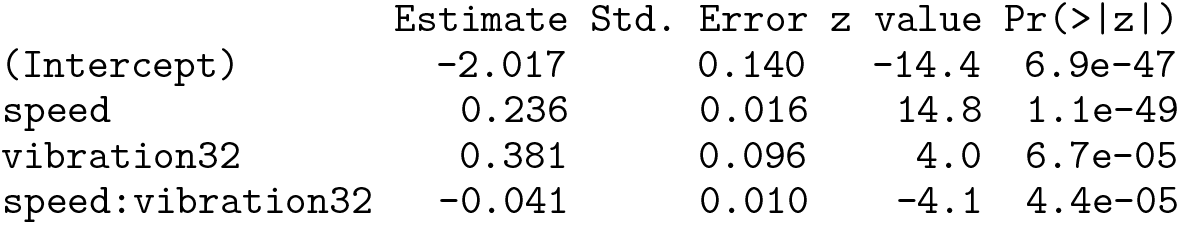

The predictors in the first and second row of the model summary correspond to *β*_0_ and *β*_1_ in Eq. 5. That is, they are intercept and slope of the response in the baseline condition, i.e. when `vibration` has a value of zero. The third and fourth rows account for the *difference* –with respect to the baseline– in intercept and slope associated with the second level of the categorical predictor (`vibration = 32`). In other words, the categorical variable `vibration` is treated by default as a “dummy variable” (James et al. 2013). In order to obtain an estimate of the coefficients associated with the second level of the categorical predictor (i.e. *β*_2_ and *β*_3_ in Eq. 5), we need to compute the algebraic sum between the dummy-coded coefficients and the corresponding baseline coefficients.

Like for the univariable GLMM, we can plot the model (Fig. 3) and obtain a fast estimate based on delta method of the psychometric parameters with **MixedPsy** functions:

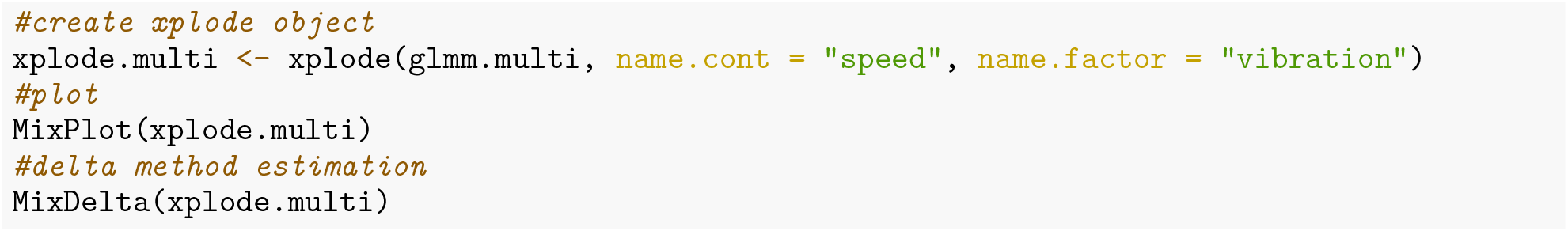

**Figure 3:**
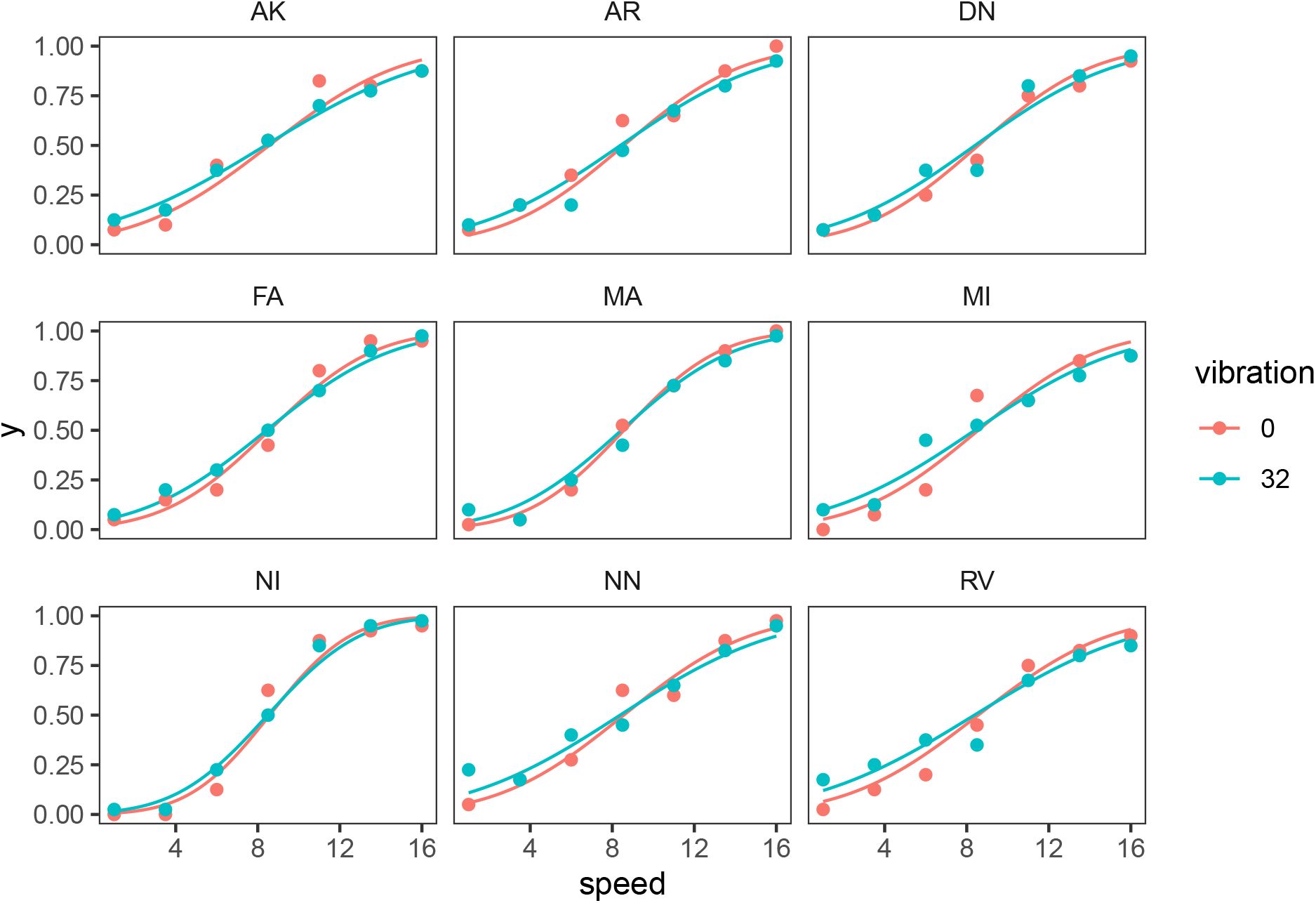
Plot of the multivariable GLMM using `MixPlot()`. By default, the function provides a different panel for each participant. To obtain panels diveded by levels of the catecorical variable, use argument `facet_by`.

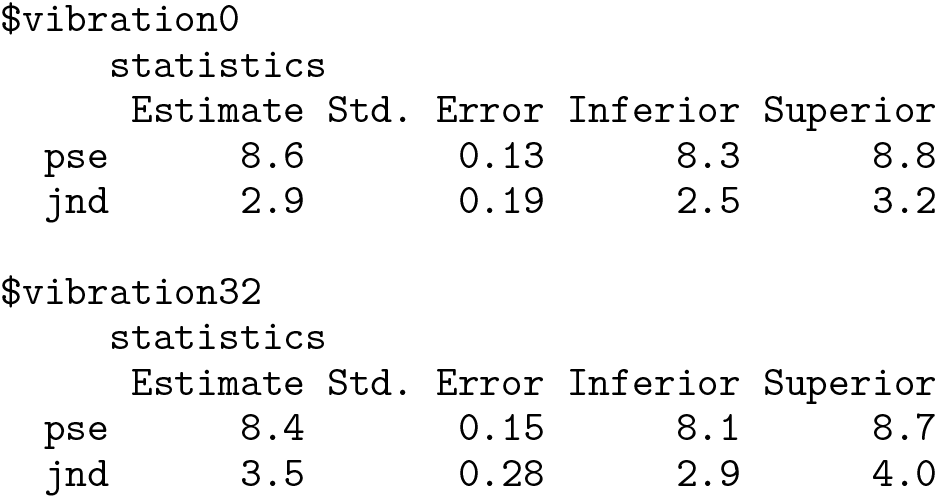

To calculate the bootstrap-based estimate of the psychophysical parameters, we use `pseMer()`. For a multivariable GLMM, we must specify how to combine the predictors of the linear model to estimate the psychometric parameters, taking into account the equations that relate PSE and JND to the GLMM coefficients (Eq. 2 and 3), as well as the fact that such coefficients are expressed with dummy coding:

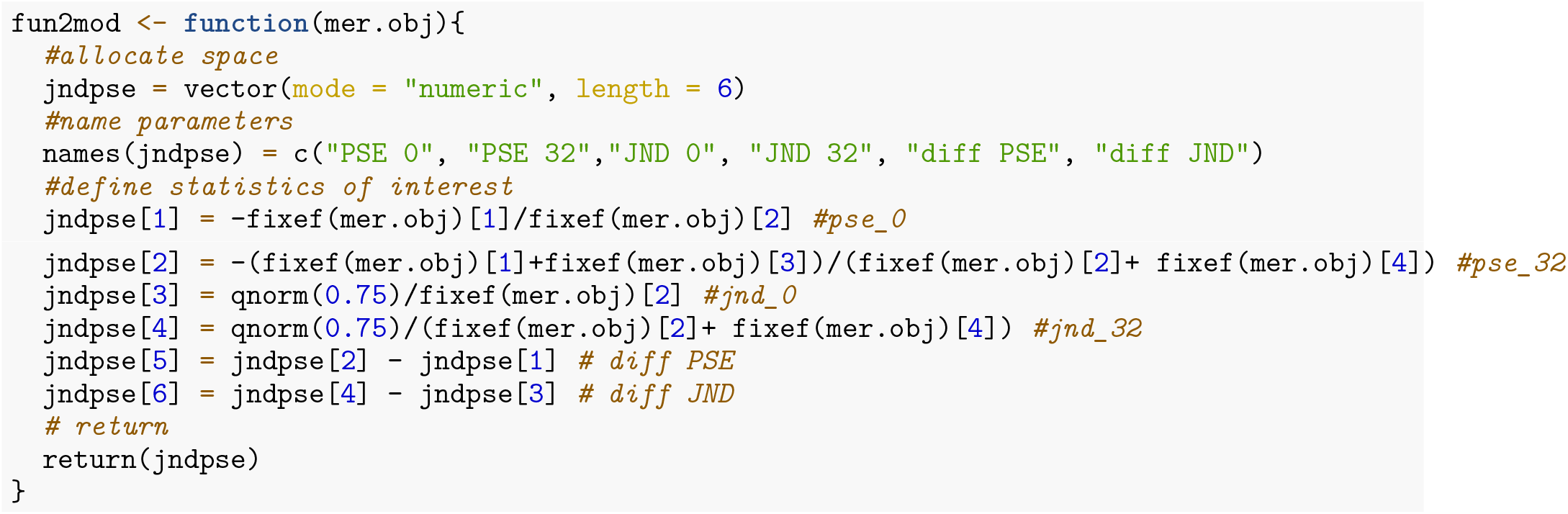

The fixed-effects estimates in an object of class `glmerMod` are extracted with function `fixef()`. The number in square brackets is the index of the specific estimate, following the order displayed in the model summary. In the case of our example model `glmm.multi`, the first and second estimate are intercept and slope in the baseline condition (no masking vibration), respectively. The third and fourth are the difference in intercept and slope associated with the change in vibration (32 Hz vibration). In our function, we considered 6 parameters of interest. Parameters 1 and 2 (`jndpse[1]` and `jndpse[2]` in the `fun2mod()`) provide estimates for PSE in the baseline and experimental condition, respectively. According to Eq. 2 and Eq. 5, PSE calculated for the two conditions of the categorical variable included in the model have the form: *PSE*_1_ = –*β*_0_/*β*_1_, *PSE*_2_ = –*β*_2_/*β*_3_. Note that, since the model defined with `glmer()` uses dummy coding, the PSE for the second condition is calculated from the algebraic sum of the fixed-effects.Analogous considerations apply for the JND estimates defined in parameters `jndpse[3]` and `jndpse[4]`. With the last two parameters, we estimate how PSE and JND differ between the two conditions. The function is then provided as input argument to `pseMer()` for the bootstrap simulation:

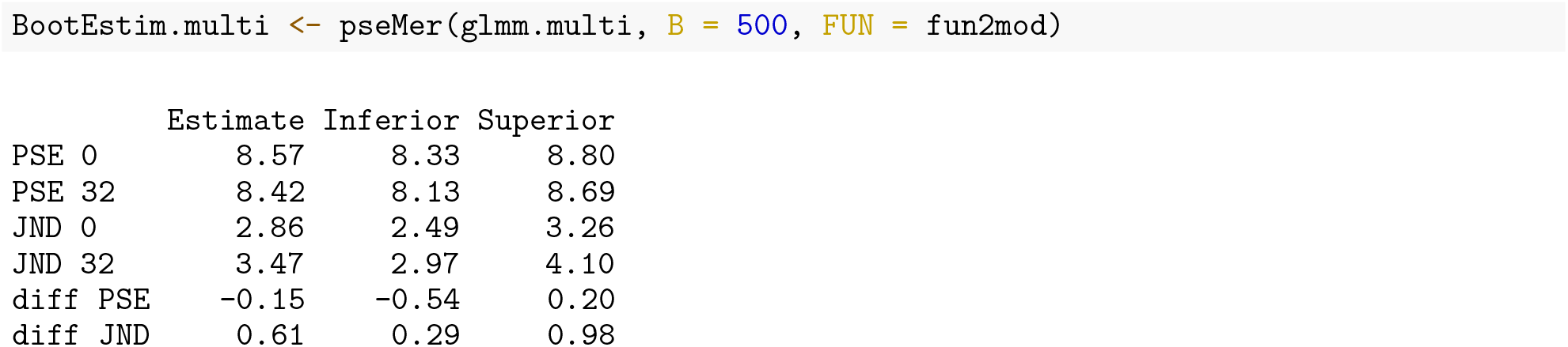

This bootstrap estimation took 3.92 minutes to be completed with our computer. When considering the psychometric parameters, the bootstrap estimations are consistent, although not identical, with those obtained with `MixDelta()`. Other relevant information can be inferred by evaluating the estimates of the differences in PSE and JND between experimental conditions: namely, the 95% confidence interval of the PSE difference includes 0, indicating that we cannot reject the null hypothesis for which the PSEs do not differ between the conditions. On the contrary, the confidence interval of the JND difference lies above zero, indicating a significant effect of the masking vibration on this parameter. These results are consistent with those reported in (Dallmann, Ernst, and Moscatelli 2015), Experiment 1b.

## Discussion

In this tutorial, we provided a brief overview of the theoretical and methodological background for collecting psychophysical data, and showed two exemplary pipelines for analyzing the resulting dataset: the first involved fitting GLMs to the response of individual participants, followed by second-level analysis on the obtained psychometric parameters (i.e. their PSEs and JNDs). The other consisted in evaluating the effects of the experimental predictors on the response by fitting a GLMM to the entire dataset, and estimate PSE and JND across all participants (conditional estimate.) While both approaches are equally valid and easy to implement in R, we showed in our previous study (Moscatelli, Mezzetti, and Lacquaniti 2012) that the GLMM framework offers several advantages over the more classical two-level analysis, such as higher statistical power and easier assessment of the goodness of fit of the model. Our package **[MixedPsy](https://cran.r-project.org/package=MixedPsy)** provides a set of tools for facilitating the analysis of psychophysical data with both GLMs and GLMMs in R. By providing ways to easily implement both approaches within the same package, and by introducing basic ideas in this tutorial in a non-technical fashion, we hope to make the GLMM framework accessible also for researchers that are less familiar with the related statistical methods for psychophysics.

In addition to providing functions for estimating psychophysical parameters, our package allows for a straightforward visualization of the fitted GLMs and GLMMs with functions `PsychPlot()` and `MixPlot()`, respectively. Model visualization is a fundamental part of the analysis pipeline, both in an exploratory phase (Wickham and Grolemund 2016) as well as when communicating results. While being useful for a quick inspection of the fitted models, the functions in **MixedPsy** do not allow for a fine control of the plot aesthetics, which is often necessary to accurately represent the information contained in the data. In order to facilitate model visualization and exploratory data analysis, we provided some source code examples for creating figures with **ggplot2** in the Appendix. For further information on these topics, refer to (Wickham 2016; Wickham and Grolemund 2016).

It is important to remember that GLMMs do not rely on statistical assumptions of asymptotic normal distribution (Agresti 2002; Casella and Berger 2002). For this reason, it is generally recommended to use simulations rather than analytic methods for variance estimation of PSEs and JNDs obtained from GLMM parameters (James et al. 2013). In **MixedPsy**, both methods are available, i.e. bootstrap simulation with `pseMer()` and delta-method estimation with `MixDelta()`. We suggest to use the latter for a fast evaluation of the effects in the phase of exploratory data analysis, but confirm the estimation with the bootstrap method when reporting results.

Simulation methods are also recommended for power analysis of mixed models. We refer interested readers to R package **[simr](https://cran.r-project.org/package=simr)** (Green and MacLeod 2016), which provides tools based on Monte Carlo simulations to calculate power for GLMMs fitted with **lme4**, and to tutorials dedicated specifically to the topic (Brysbaert and Stevens 2018; Kumle, Võ, and Draschkow 2021).

There are many other excellent packages in R dedicated to providing tools for psychophysical data, which are available on CRAN and listed in the [CRAN Task View: Psychometric Models and Methods](https://cran.r-project.org/web/views/Psychometrics.html). In particular, we refer to packages such as **[psyphy](https://cran.r-project.org/package=psyphy)** (Knoblauch 2022), **[quickpsy](https://cran.r-project.org/package=quickpsy)** (Linares, López-Moliner, et al. 2016), **[MPDiR](https://cran.r-project.org/package=MPDiR)** (Knoblauch and Maloney 2021), for fitting psychometric functions for the analysis of the responses of individual participants. While **MixedPsy** and the packages mentioned above use frequentist approaches to estimate psychometric parameters, package **[MCMCglmm](https://cran.r-project.org/package=MCMCglmm)** (Hadfield 2010) uses Bayesian inference to extract the information contained in experimental data, by means of Markov Chains Monte Carlo techniques. Bayesian inference has been validated for the analysis of psychophysical data (Kuss, Jäkel, and Wichmann 2005). Previous knowledge from the researcher on the distribution of a given parameter (for example based on the literature or on a preliminary experiment) can be included in the model in form of a prior distribution.

Other free and open-source tools for analyzing psychophysical data are implemented in popular programming environments such as Matlab and Python. In Matlab, [Palmedes](https://www.palamedestoolbox.org/) (Prins and Kingdom 2018) is a relatively new toolbox for psychophysical research, that allows users to fit psychometric functions and perform model comparisons. **psignifit** (Schütt et al. 2016), currently on its fourth version, is a classical package for estimating the psychometric function, available for both [Matlab](https://github.com/wichmann-lab/psignifit/wiki) and [Python](https://github.com/wichmann-lab/python-psignifit/wiki). Other resources for psychophysics in Python are included in [PsychoPy](https://www.psychopy.org/), an open-source module used by a large community of researchers in neuroscience and experimental psychology. While it is mainly used for running behavioral experiments using Python and JavaScript, PsychoPy also provides basic tools for fitting and plotting psychometric functions for individual participants with module **[pylab](https://psychopy.org/coder/tutorial3.html)**. Similarly to **lme4** in R, Python module **[Statsmodel](https://www.statsmodels.org/)** (Seabold and Perktold 2010) allows for the estimation of several statistical models, including GLMMs. **[gpboost](https://github.com/fabsig/GPBoost)** is a library for fast and reliable estimate of random effect models (Sigrist 2020). It is written in C++ and can be operated either from [R](https://github.com/fabsig/GPBoost/tree/master/R-package) or [Python](https://github.com/fabsig/GPBoost/tree/master/python-package) with their dedicated packages.

While programming environments such as R, Python, and Matlab are well established and widely used in the research community, [Julia](https://julialang.org/) is a relatively new language, that has the merit of being very fast and easy to implement. Specific routines for fitting models in psychophysics are not available in Julia, however it is possible to fit both GLMs and GLMMs with package [juliastats](https://juliastats.org/) and MixedModels.jl, respectively.

As can be seen from this list, several powerful tools are available, implementing statistical methods for psychophysical experiments. We hope that this provides orientation for navigating these tools.

## Acknowledgments

We would like to thank Prof. Maura Mezzetti and Prof. Francesco Lacquaniti for useful comments and suggestions. Thanks to Chris Geekie for the helpful comments on the language of the text. Funding: Author PB and ME were funded by the Deutsche Forschungsgemeinschaft (DFG, German Research Foundation) – Projektnummer 251654672 – TRR 161. Author AM was funded by the Italian Ministry of Health (IRCCS Fondazione Santa Lucia, Ricerca Corrente); by the Italian Ministry of Education and Research (MIUR) in the framework of PRIN 2017 (Programmi di Ricerca Scientifica di Rilevante Interesse Nazionale) with project Grant number 2017SB48FP.

## Appendix

### Basic information on R and relevant R packages

While an exhaustive overview of the R programming language is out of the scope of this tutorial, we hope that the provided information can be easily followed by users without previous or little coding experience. For more resources on the topic, we suggest to choose among the many sources of freely available material for learning R, as well as to rely on the very active community of R users (https://stackoverflow.com/tags/r/info).

A popular and user-friendly IDE choice for R is RStudio^1^, which allows to keep console and editor in the same environment while also offering additional features (e.g. syntax highlighting, tools for plotting, debugging, environment space and history management).

R packages provide functions and dataset developed by the user community that extend the basic R functionalities, freely available in R project’s software repository (CRAN) and on GitHub. Packages in CRAN can be installed using the command `install.packages(“<package-name>”)`, and loaded in the ongoing R session with `library(“<package-name>”)`. The description of a package can be accessed with command `help(package = “<package-name>”)`, the documentation of a function with `?<function-name>`.

Our package **MixedPsy** is intended as a tool to facilitate modeling and visualization of psychophysical data in R, both at a single-subject and at a population level. It is open source and can be downloaded from CRAN (stable release) and [GitHub](https://github.com/moskante/__MixedPsy__) (stable release and development version).

Routines for fitting GLMMs in **MixedPsy** are partially based and dependent on the **lme4** package (Bates et al. 2015), which provides functions for fitting and analyzing linear, generalized linear, and nonlinear mixed models.

Functions for plotting fitted models in **MixedPsy** consist in wrappers of the **ggplot2** package (Wickham 2016), which is a system for creating publication-quality graphics in R by combining instructions for mapping data variables to different aesthetics.

**ggplot2** and functions mentioned throughout the tutorial are included in **tidyverse** (Wickham et al. 2019), a collection of different packages designed for data science (Wickham and Grolemund 2016).

### Setup

The examples proposed in this tutorial were tested in the environment described below:

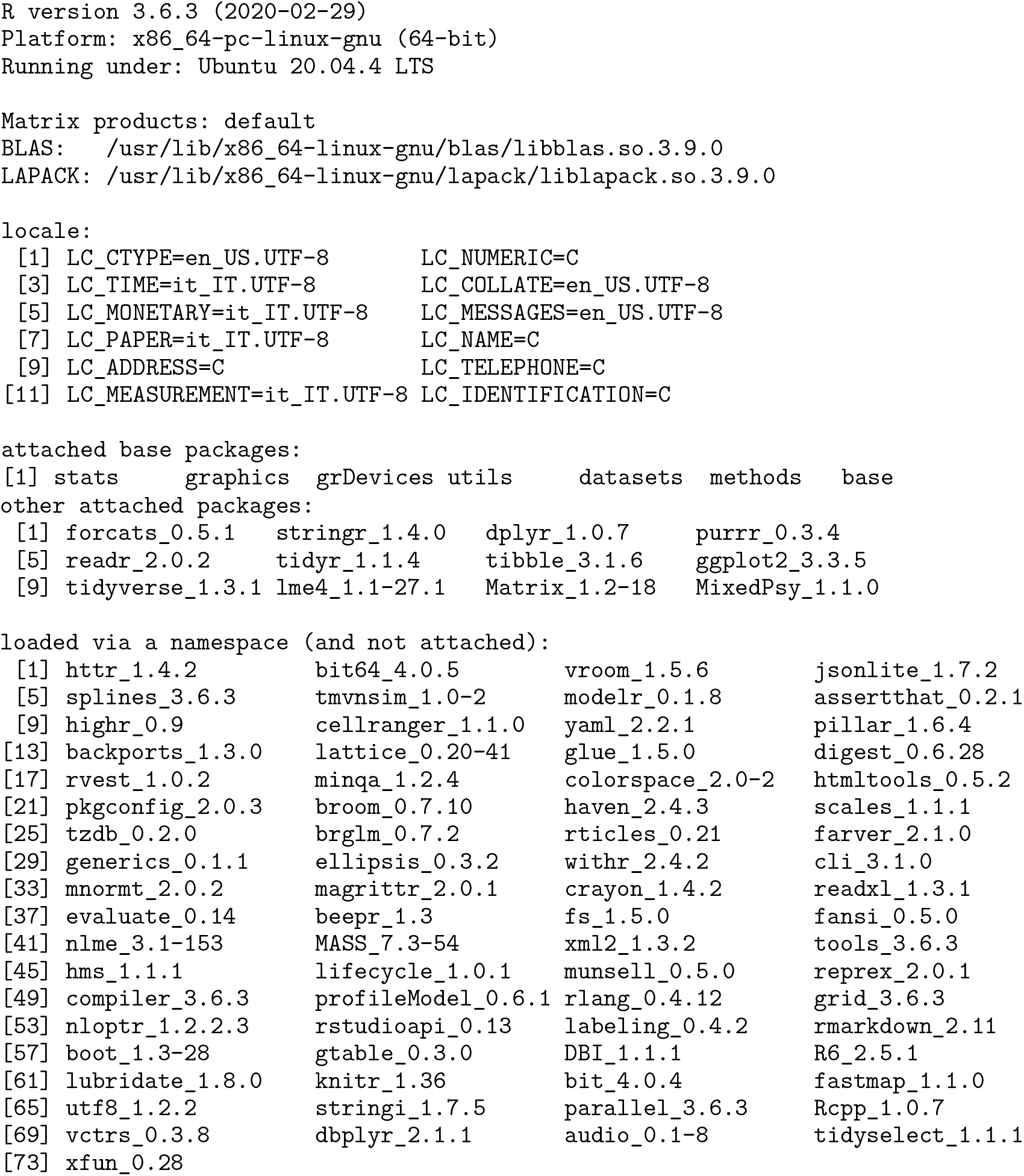

### Data visualization

Function `geom_smooth()` allows to add smoothed means and regression lines in a `ggplot` object, and it is a powerful way to show patterns in the data during exploratory analysis.

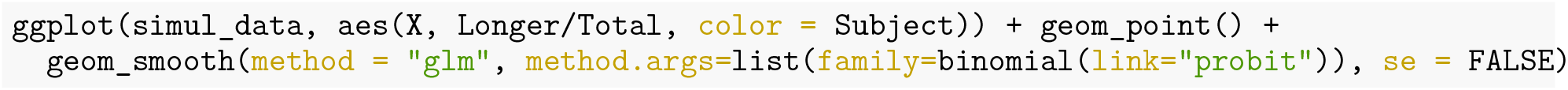

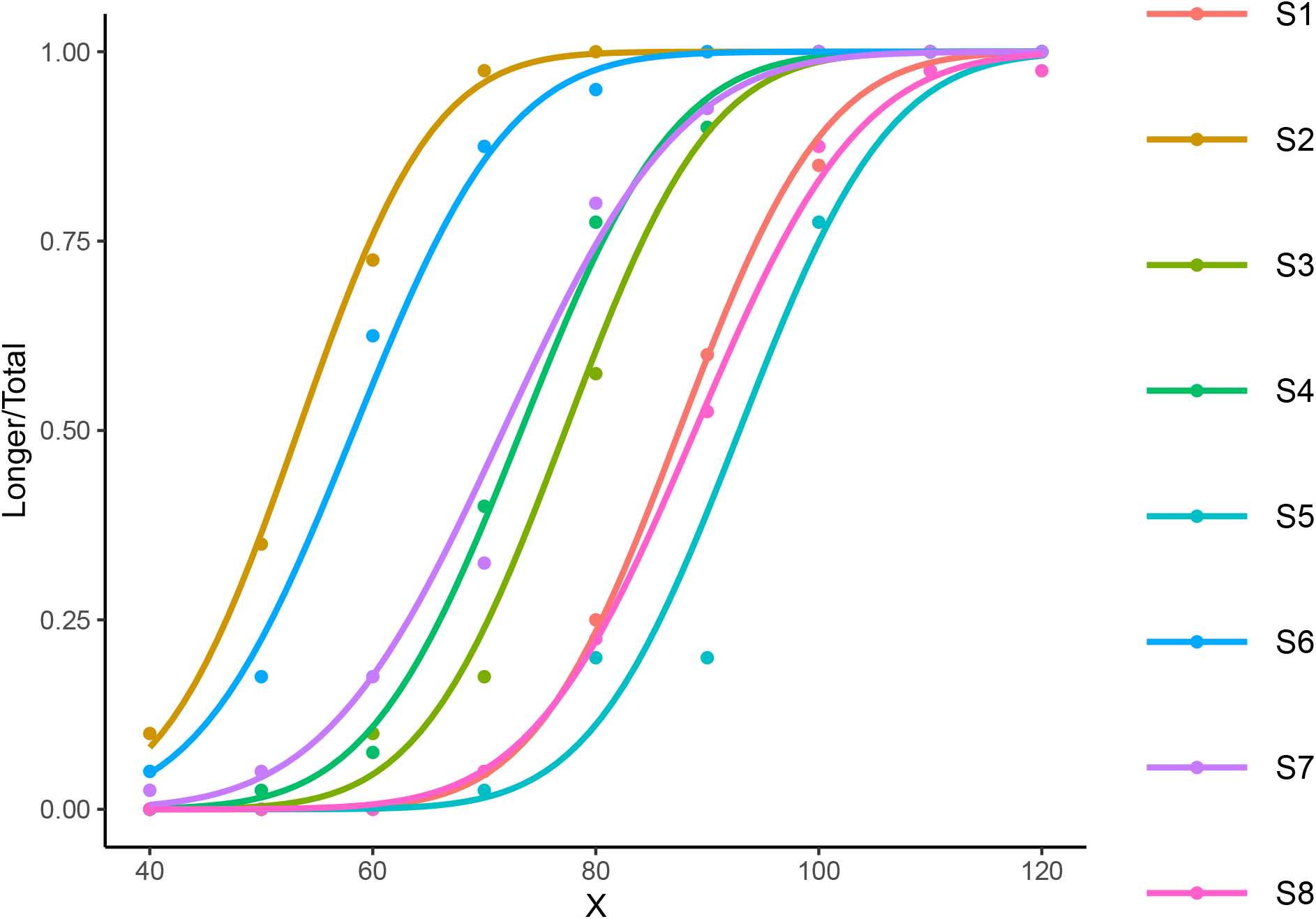

`predict()` generates numerical predictions of the fitted models, which can then be rendered in plots. In order to plot non-linear models with a smooth line and avoid to display individual segments, it is often necessary to display the model predictions using a finer mesh for the continuous variable. To do so, an option is to create a new data frame with function `expand.grid()`, and use a method for interpolating the intensity values of the continuous predictors in the range of the presented stimulus intensities. A way to do so is with function `pretty()`.

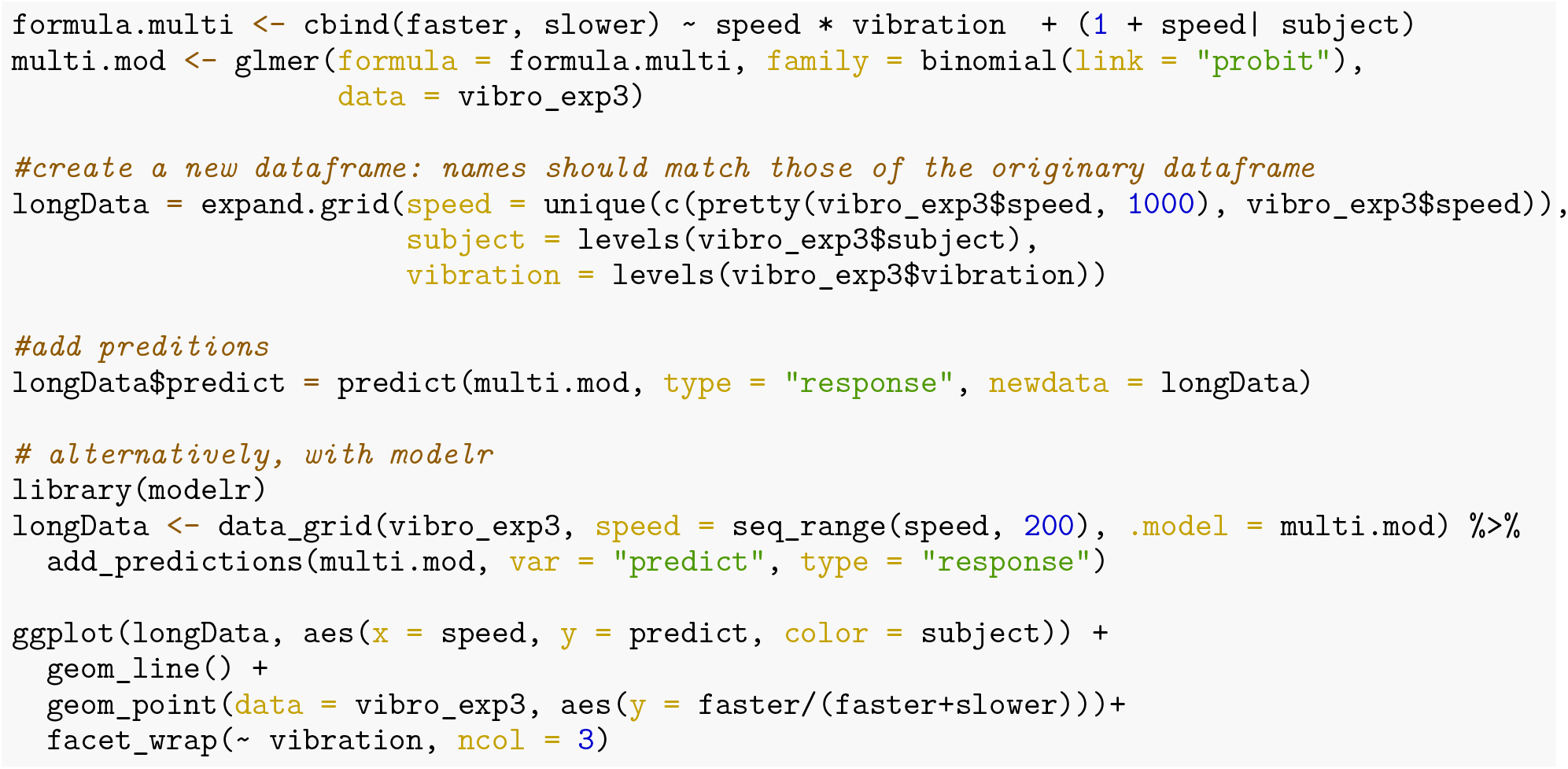

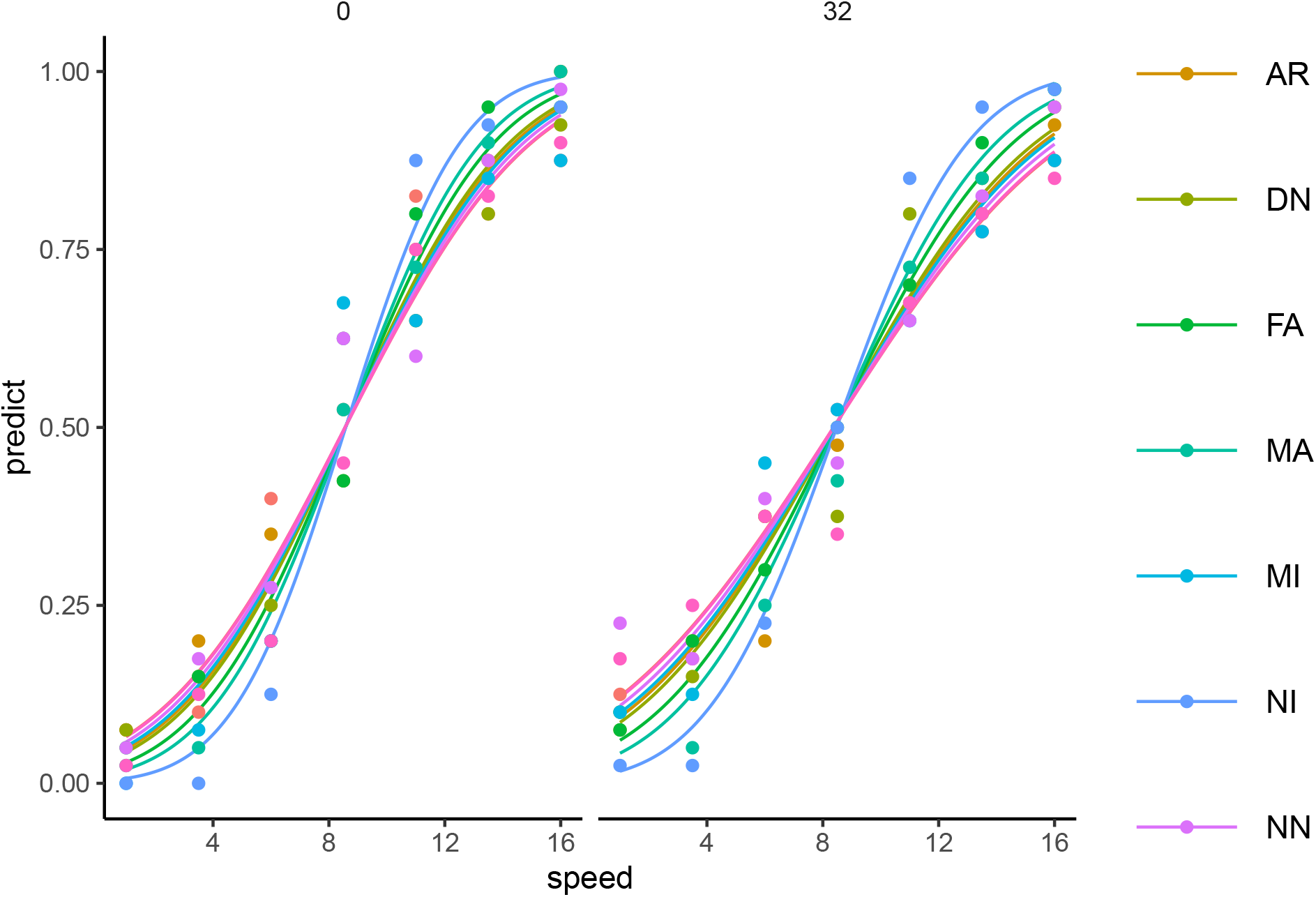

## Supplemental Data

The example dataset `exampledata.csv` and the source code of the examples illustrated in the tutorial `tutorialMixedPsy.R` are available for download from the GitHub repository MixedPsy_companion_data.

## Footnotes

1 https://rstudio.com

## Notes

https://github.com/moskante/MixedPsy_companion_data

### Competing Interest Statement

The authors have declared no competing interest.

## References

Agresti, Alan. 2002. Categorical Data Analysis. Vol. 482. John Wiley & Sons.

Akaike, Hirotugu. 1973. “Information Theory and an Extension of the Maximum Likelihood Principle,[w:] Proceedings of the 2nd International Symposium on Information, Bn Petrow, f.” Czaki, Akademiai Kiado, Budapest.

Bates, Douglas, Martin Mächler, Ben Bolker, and Steve Walker. 2015. “Fitting Linear Mixed-Effects Models Using lme4.” Journal of Statistical Software 67 (1): 51. https://doi.org/10.18637/jss.v067.i01.

Brysbaert, Marc, and Michaël Stevens. 2018. “Power Analysis and Effect Size in Mixed Effects Models: A Tutorial.” Journal of Cognition 1 (1).

Casella, George, and Roger L Berger. 2002. Statistical Inference. Vol. 2. Duxbury Pacific Grove, CA.

Dallmann, Chris J., Marc O. Ernst, and Alessandro Moscatelli. 2015. “The role of vibration in tactile speed perception.” Journal of Neurophysiology 114 (6): 3131–39. https://doi.org/10.1152/jn.00621.2015.

Green, Peter, and Catriona J. MacLeod. 2016. “Simr: An r Package for Power Analysis of Generalised Linear Mixed Models by Simulation.” Methods in Ecology and Evolution 7 (4): 493–98. https://doi.org/10.1111/2041-210X.12504.

Hadfield, Jarrod D. 2010. “MCMC Methods for Multi-Response Generalized Linear Mixed Models: The MCMCglmm R Package.” Journal of Statistical Software 33 (2): 1–22. https://www.jstatsoft.org/v33/i02/.

Hidalgo, Bertha, and Melody Goodman. 2013. “Multivariate or multivariable regression?” American Journal of Public Health 103 (1): 39–40. https://doi.org/10.2105/AJPH.2012.300897.

James, Gareth, Daniela Witten, Trevor Hastie, and Robert Tibshirani. 2013. An Introduction to Statistical Learning. Vol. 112. Springer.

Kingdom, Frederick A. A., and Nicolaas and Prins. 2016. Psychophysics: A Practical Introduction, 2nd Edn. Academic Press.

Klein, Stanley A. 2001. “Measuring, Estimating, and Understanding the Psychometric Function: A Commentary.” Perception & Psychophysics 63 (8): 1421–55.

Knoblauch, Kenneth. 2022. Psyphy: Functions for Analyzing Psychophysical Data in r. https://CRAN.R-project.org/package=psyphy.

Knoblauch, Kenneth, and Laurence T. Maloney. 2012. Modeling Psychophysical Data in R. New York, NY: Springer New York. https://doi.org/10.1007/978-1-4614-4475-6.

Knoblauch, Kenneth, and Laurence T. Maloney. 2021. MPDiR: Data Sets and Scripts for Modeling Psychophysical Data in r. https://CRAN.R-project.org/package=MPDiR.

Kumle, Leah, Melissa L-H Võ, and Dejan Draschkow. 2021. “Estimating Power in (Generalized) Linear Mixed Models: An Open Introduction and Tutorial in r.” Behavior Research Methods 53 (6): 2528–43.

Kuss, Malte, Frank Jäkel, and Felix A. Wichmann. 2005. “Bayesian inference for psychometric functions.” Journal of Vision 5 (5): 8–8. https://doi.org/10.1167/5.5.8.

Linares, Daniel, Joan López-Moliner, et al. 2016. “Quickpsy: An r Package to Fit Psychometric Functions for Multiple Groups.” The R Journal 8 (1): 122–31.

Moscatelli, Alessandro, and Priscilla Balestrucci. 2021. Psychophysics with r: The r Package MixedPsy. https://CRAN.R-project.org/package=MixedPsy.

Moscatelli, Alessandro, M. Mezzetti, and Francesco Lacquaniti. 2012. “Modeling psychophysical data at the population-level: The generalized linear mixed model.” Journal of Vision 12 (11): 26. https://doi.org/10.1167/12.11.26.

Pelli, Denis G, and Farell Bart. 1991. “Psychophysical Methods.” In Handbook of Optics, 2nd Ed., 29. 21–13.

Prins, Nicolaas, and Frederick A. A. Kingdom. 2018. “Applying the Model-Comparison Approach to Test Specific Research Hypotheses in Psychophysical Research Using the Palamedes Toolbox.” Frontiers in Psychology 9. https://doi.org/10.3389/fpsyg.2018.01250.

R Core Team. 2020. R: A Language and Environment for Statistical Computing. Vienna, Austria: R Foundation for Statistical Computing. https://www.R-project.org/.

Rohde, Marieke, Loes CJ van Dam, and Marc O Ernst. 2016. “Statistically Optimal Multisensory Cue Integration: A Practical Tutorial.” Multisensory Research 29 (4-5): 279–317.

Schütt, Heiko H., Stefan Harmeling, Jakob H. Macke, and Felix A. Wichmann. 2016. “Painfree and Accurate Bayesian Estimation of Psychometric Functions for (Potentially) Overdispersed Data.” Vision Research 122: 105–23. https://doi.org/10.1016/j.visres.2016.02.002.

Seabold, Skipper, and Josef Perktold. 2010. “Statsmodels: Econometric and Statistical Modeling with Python.” In 9th Python in Science Conference.

Sigrist, Fabio. 2020. “Gaussian Process Boosting.” arXiv Preprint arXiv:2004.02653.

Wichmann, Felix A., and N. Jeremy Hill. 2001. “The psychometric function: I. Fitting, sampling, and goodness of fit.” Perception and Psychophysics 63 (8): 1293–313. https://doi.org/10.3758/BF03194544.

Wickham, Hadley. 2016. Ggplot2: Elegant Graphics for Data Analysis. Springer-Verlag New York. https://ggplot2.tidyverse.org.

Wickham, Hadley, Mara Averick, Jennifer Bryan, Winston Chang, Lucy D’Agostino McGowan, Romain François, Garrett Grolemund, et al. 2019. “Welcome to the Tidyverse.” Journal of Open Source Software 4 (43): 1686.

Wickham, Hadley, and Garrett Grolemund. 2016. R for Data Science: Import, Tidy, Transform, Visualize, and Model Data. O’Reilly Media, Inc. https://r4ds.had.co.nz.

Yssaad-Fesselier, Rosa, and Kenneth Knoblauch. 2006. “Modeling Psychometric Functions in r.” Behavior Research Methods 38 (1): 28–41.

